# Machine learning of cellular metabolic rewiring

**DOI:** 10.1101/2023.08.11.552957

**Authors:** Joao B. Xavier

## Abstract

Metabolic rewiring allows cells to adapt their metabolism in response to evolving environmental conditions. Traditional metabolomics techniques, whether targeted or untargeted, often struggle to interpret these adaptive shifts. Here, we introduce *MetaboLiteLearner*, a machine learning framework that harnesses the detailed fragmentation patterns from electron ionization (EI) collected in scan mode during gas chromatography/mass spectrometry (GC/MS) to predict abundance changes in metabolically adapted cells. When tested on breast cancer cells with different preferences to metastasize to specific organs, *MetaboLiteLearner* predicted the impact of metabolic rewiring on metabolites withheld from the training dataset using only the EI spectra, without metabolite identification or pre-existing knowledge of metabolic networks. The model learned captures shared and unique metabolomic shifts between brain- and lung-homing metastatic lineages, suggesting potential organ-tailored cellular adaptations. Integrating machine learning and metabolomics paves the way for new insights into complex cellular adaptations.

**Significance:** Metabolic rewiring—the cellular adaptation to shifts in environment and nutrients—plays key roles in many contexts, including cancer metastasis. Traditional metabolomics often falls short of capturing the nuances of these metabolic shifts. This work introduces *MetaboLiteLearner*, a machine learning approach that harnesses the rich fragmentation patterns from electron ionization collected in scan mode during gas chromatography/mass spectrometry, paving the way for new insights into metabolic adaptations. Demonstrating its robustness on a breast cancer model, we highlight *MetaboLiteLearner*’s potential to reshape our understanding of metabolic rewiring, with implications in diagnostics, therapeutics, and basic cell biology.

## Introduction

Cells dynamically rewire their intracellular metabolism in response to changing nutrients, signals, and environmental cues (1). These adjustments allow cells to maintain homeostasis, optimize energy production, and fulfill the biosynthetic demands for growth and repair. From unicellular organisms like yeast adapting to changing nutrient availability (2) to the cells of multicellular organisms during development (3), disease (4), or environmental stress (5), metabolic rewiring is a universal feature of cellular adaptation and survival.

Metabolic rewiring shifts the biochemical composition of cells, and the changes observed in the intracellular metabolome provide a window into the underlying cellular adaptation (6). Traditional approaches to metabolomics are either targeted or untargeted. Targeted metabolomics allows for the accurate and sensitive quantification of a predefined set of metabolites but risks missing novel or unexpected ones (7, 8). Untargeted metabolomics allows for the comprehensive survey of all detectable metabolites within a biological sample but has challenges regarding reproducibility and identifying unknown compounds (9).

To address these limitations, we present *MetaboLiteLearner*. This novel computational method uses machine learning to investigate metabolic rewiring using metabolomic data without relying on prior knowledge of the metabolic network or having to identify the compounds from their spectra. *MetaboLiteLearner* deploys the extensive molecular fragmentation achieved through electron ionization (EI) in gas chromatography/mass spectrometry (GC/MS) and acquired in scan mode. The fragmentation patterns are treated as input features in a supervised learning model, which associates these features with the changes in metabolite abundance observed in cells undergoing metabolic rewiring. *MetaboLiteLearner* effectively uses information about the molecular structure of metabolites to predict how their abundances change in cells adapted metabolically to diverse circumstances.

As proof of concept, we utilized clones of the MDA-MB-231 breast cancer cell line that home to the lung and brain (10). These derivatives, developed through *in vivo* selection in mice (11, 12), show pronounced transcriptomic shifts in laboratory cultures. In a previous study, we employed targeted metabolomics to analyze a comprehensive panel of 645 metabolites from these cells. That analysis unveiled distinct intracellular metabolic profiles between the brain- and lung-homing cells and their parental counterparts—distinct metabolic rewiring patterns that were posteriorly validated through direct metabolic exchange measurements and indirect assessments via stable isotope tracing (13). Given this detailed characterization, the MDA-MB-231 system serves as an exemplary model to benchmark the efficacy of *MetaboLiteLearner*.

*MetaboLiteLearner* identified unique metabolic features in brain- and lung-homing cells that may be adaptations to the specific challenges of their target organs. This insight aligns with the findings from our earlier study (13), which relied on a large panel of metabolites validated and measured through LC/MS, a method considered more laborious and expensive. On the other hand, *MetaboLiteLearner* draws insights using all the data produced from GC/MS in scan mode, without relying on prior information such as spectral libraries or extensive validation to draw its insights. *MetaboLiteLearner* can be applied to various cell types and conditions, allowing researchers in different fields to explore new aspects of cellular metabolism. Beyond streamlining the analytical process, *MetaboLiteLearner* paves the way for in-depth insights into metabolic rewiring patterns by finding associations between molecular structure and metabolic adaptability.

## Results

### *MetaboLiteLearner*: Theory and Application

*MetaboLiteLearner* capitalizes on GC/MS data from intracellular metabolome extracts. The extracts are primed for GC/MS analysis using trimethylsilyl (TMS) derivatization, which enhances metabolite volatility. EI fragmentation to provide a detailed molecular structure spectrum (14). The TMS derivatization, combined with GC/MS and EI fragmentation, produces reproducible spectra and retention times for each metabolite. Convetional targeted metabolomics often measures specific ion peak areas (**Fig. 1A**). Here, *MetaboLiteLearner* harnesses the entirety of data available in scan mode (**Fig. 1B**). This spans ion fragment abundances from 50 to 600 mass-to-charge ratio units (m/z) and all GC retention times.

**Figure 1.**
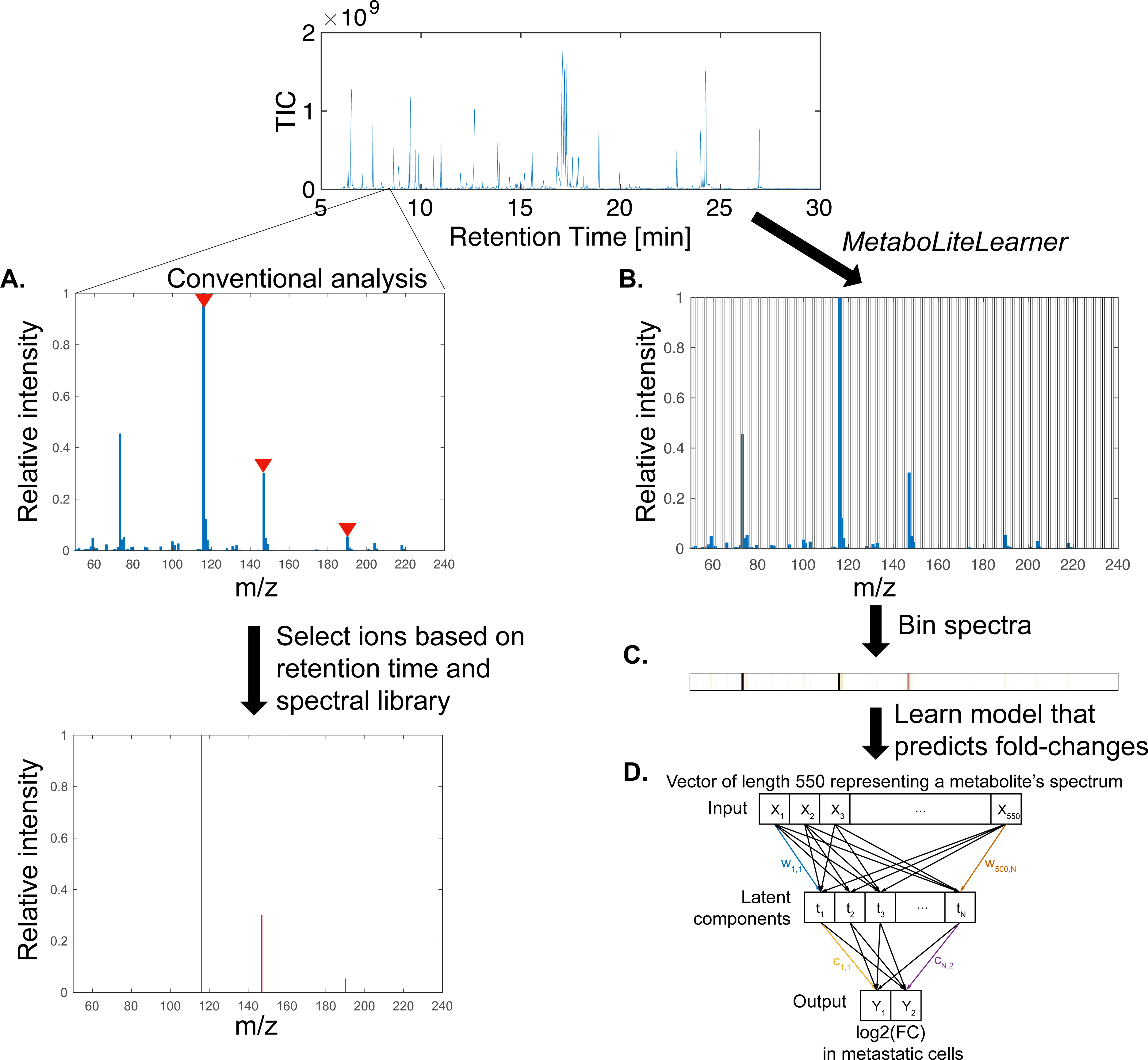
Workflow and Functionality of *MetaboLiteLearner*. **(A)** Traditional targeted metabolomics uses specific ion peak areas. **(B)** *MetaboLiteLearner* uses GC/MS data acquired in scan mode in *MetaboLiteLearner*, which encompasses ion fragment abundances ranging from 50 to 600 m/z captured at all GC retention times. **(C)** Each molecule’s electron impact (EI) fragmentation spectrum is depicted as a 550-dimensional vector. These high-dimensional vectors are paired with their corresponding log2 fold change values, which serve as training labels. **(D)** The transformation matrices (‘w’ for spectra and ‘c’ for log2 fold changes) are the Partial Least Squares Regression (PLSR) loadings used to map data into the *N*-dimensional latent space. This enables *MetaboLiteLearner* to learn the relationship between metabolite structure and metabolic rewiring.

*MetaboLiteLearner* integrates full spectra with the corresponding log two-fold changes in abundance, reflecting their abundance changes due to metabolic rewiring. Each molecule’s EI fragmentation spectrum is encoded as a 550-dimensional vector (**Fig. 1C**), with corresponding log2 fold changes (obtained by comparing baseline vs. rewired cell metabolite abundances) as the training data labels. Following the supervised learning paradigm, the model’s efficacy is gauged by predicting log2 fold changes on a new compound set and comparing these predictions with actual data (15).

*MetaboLiteLearner* uses Partial Least Squares Regression (PLSR) to construct a linear model by projecting the predictors and the response variables onto a new *N*-dimensional space (16). PLSR generates a model that balances complexity and interpretability without sacrificing the power of linear combinations of original variables. This approach is analogous to an Artificial Neural Network (ANN) that employs a hidden layer with *N* neurons, except that PLSR employs linear functions instead of non-linear activation functions networks (17).

To transform the high-dimensional input (a 550-dimensional EI spectrum) and output (log2 fold changes) into a latent space, *MetaboLiteLearner* uses two matrices: ‘*w*’ for the spectra and ‘*c*’ for the log2 fold changes (**Fig. 1D**). The latent space dimensionality, *N*, dictates the model’s complexity. An optimal number of dimensions, *N*_*opt*_, is determined through cross-validation to prevent under- and overfitting (15). When applied to metabolomics data, the loadings derived from PLSR—coefficients that describe the relationship between the original predictors and the new latent factors—provide insights into the underlying cellular adaptations. These loadings are directly tied to the EI-fragmentation spectra of metabolites and indicate molecular structural features linked to their abundance changes in metabolically rewired cells. Therefore, once the model is trained, the loadings provide insights into the relationships between metabolite structure and metabolic rewiring, shedding light on cellular adaptation mechanisms

### Breast Cancer Cell Data Integration into *MetaboLiteLearner*

We collected the data for *MetaboLiteLearner* from the MDA-MB-231 breast cancer cell line and its brain and lung-targeted derivatives. The derivatives, originating from *in vivo* mouse selection (11, 12), exhibited transcriptomic differences compared to the original MDA-MB-231 cells in lab cultures. Our prior work identified marked intracellular metabolome variances between the brain- and lung-homing cells and the parent cells, indicating that cells have undergone metabolic rewiring (13).

We used the MDA-MB-231 breast cancer cell line and its specialized derivatives to feed data into MetaboLiteLearner. These specialized cells, developed from in vivo mouse selection, have different transcriptomic profiles than their parent cells (11, 12). The choice of these cells serves multiple purposes. First, our previous work with these cells has provided a wealth of detailed data, as a solid ground for validating *MetaboLiteLearner* (13). This earlier research show clear changes in the intracellular metabolome between brain- and lung-homing cells and their parent cells, indicating significant metabolic rewiring. Moreover, re-analyzing these cells with *MetaboLiteLearner* could reveal new metabolic changes and unidentified metabolites that were overlooked in the initial study. *MetaboLiteLearner* leverages alterations in all metabolites, even unidentified ones, allowing us to explore metabolic changes in areas of the metabolic network that are typically not covered by textbook metabolic models.

We cultured all three cell variants, and the metabolites were harvested during balanced growth, which is when cells divide at exponential growth and before any slowdown in growth due to confluency (**Fig. 2A**). We then extracted soluble metabolites, dried all samples and controls, and derivatized the polar metabolites using trimethylsilyl (TMS) derivatization. After collecting the data with GC/MS with EI in scan mode (18), all the samples were aligned to consistent retention (6-30 mins at 0.01 min intervals) and m/z (50 to 600 m/z at 1 m/z steps) ranges. The resulting individual data matrices were joined into a singular “virtual bulk sample” matrix (**Fig. 2B**).

**Figure 2.**
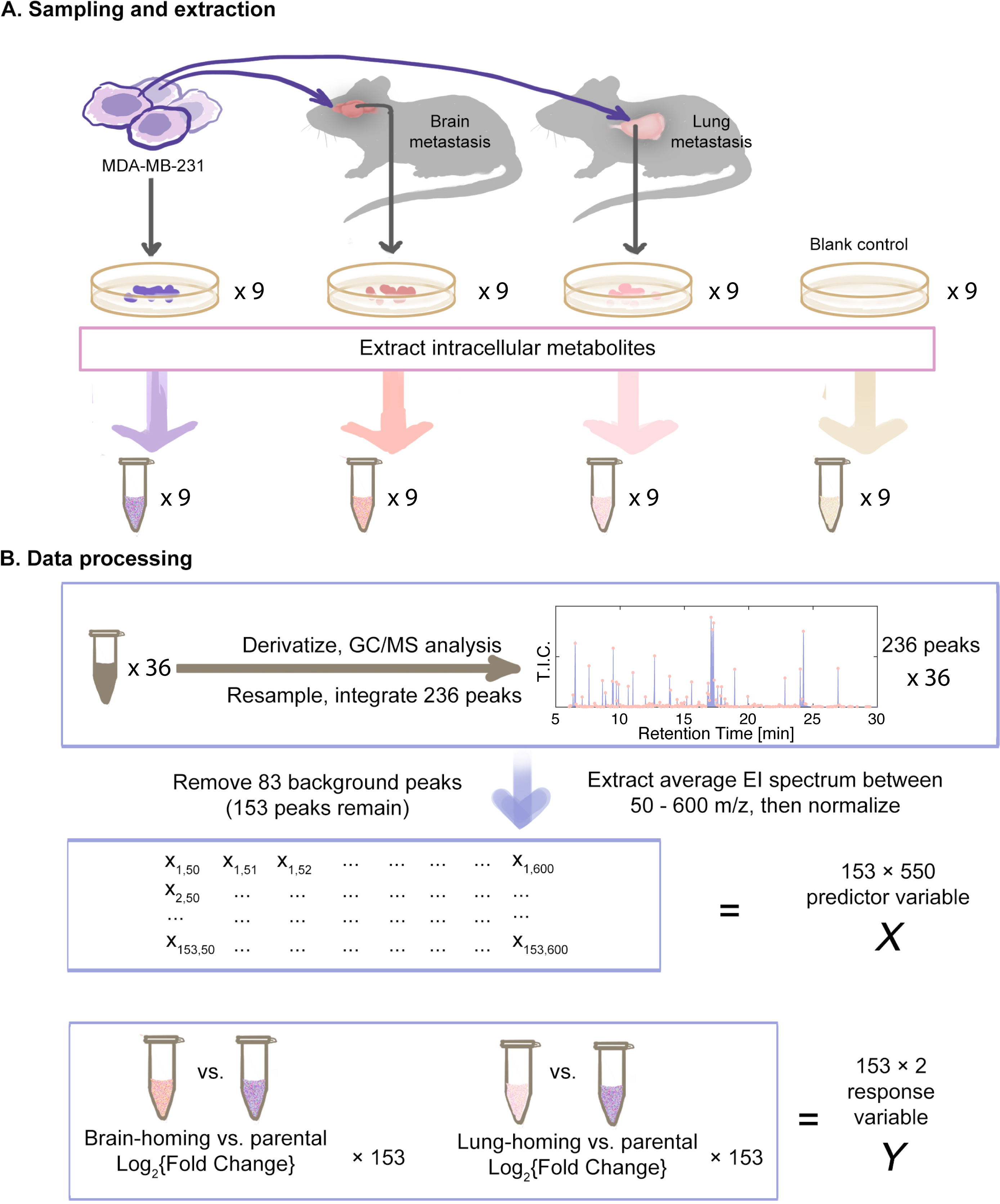
Data Acquisition and Processing for *MetaboLiteLearner* from Breast Cancer Cell Derivatives. **(A)** Cell Culture and Metabolite Extraction: Breast cancer cell lines, including the parental MDA-MB-231 cells and its brain- and lung-homing derivatives, were cultivated. These derivatives were procured through *in vivo* selection using mice. Under consistent media conditions *in vitro*, intracellular metabolites from these cells were extracted to ensure a reliable data source for subsequent processing. **(B)** GC/MS Processing and Data Aggregation: Samples underwent GC/MS analysis following the TMS derivatization protocol. The generated data matrices, unique for each sample, were amalgamated to create a virtual “bulk” sample. Peaks were identified, and their spectra were extracted from this consolidated matrix. The input (X) for *MetaboLiteLearner* encompasses the mass spectra of each peak. The output data (Y) indicates the comparative abundance shift of each peak in brain- and lung-homing cells relative to the parental cells.

Spectrum extraction from this composite data enabled us to determine the log2 fold change in metabolite abundance in the organ-homing cells relative to the parental cells through a linear mixed-effects model. Our input for *MetaboLiteLearner* consisted of spectra arrays from m/z intervals of 50 to 600, normalized to their norms. The dataset, encompassing 153 unique spectra alongside their respective log2 fold-changes, is now presented as the *MetaboLiteLearner* Open Dataset (MLOD).

### Training and Evaluating *MetaboLiteLearner*

Using the MLOD, we first determined the optimal number of latent components, *N*_*opt*_. Through hold-out cross-validation, as we increased *N* from 1 (simplest model) to 30 (most complex), the training error dropped monotonically, suggesting a better fit of the training data with increased model complexity (**Fig. 3A**). The test error reached a minimum at *N*=11 components before rising, indicating potential overfitting in the more complex models. Utilizing a conservative methodology from supervised learning (15), we selected *N*_*opt*_=5, which showcased the smallest test error within one standard error. The *N*_*opt*_=5 model predictions correlated robustly with the actual log2 fold changes (ρ=0.39, P-value≪0.01). We should note that other datasets may require a different value of *N*_*opt*_; *N*_*opt*_ should be empirically determined for every dataset.

**Figure 3.**
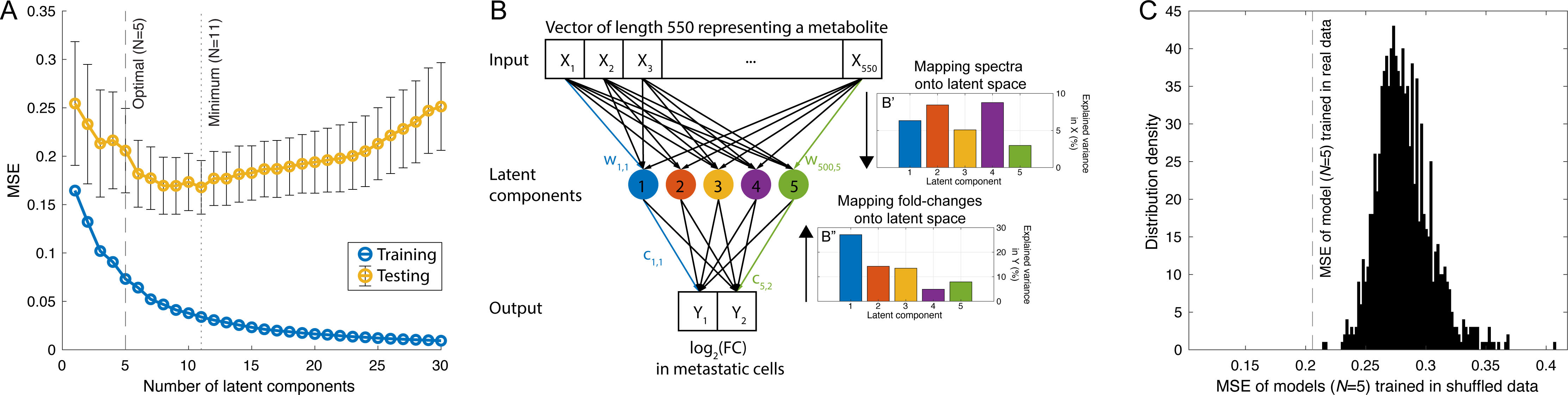
Optimization and Evaluation of MetaboLiteLearner’s Predictive Performance. **(A)** Model Complexity vs. Error: Through hold-out cross-validation, the training error consistently decreased as latent components (N) increased from 1 to 30. Test error reached its lowest at *N*=11 before it began to rise, highlighting potential overfitting with more complex models. The chosen optimal model had *N*_*opt*_=5 components, correlating strongly with true log2 fold changes (ρ=0.39, P-value≪0.01). (**B)** Schematic of the model trained with 5 components. **(B’ and B”)** Variance Explained by Latent Factors: Using the Nopt=5 model, the transformations into the 5-dimensional latent space covered 32% of the predictor variance and 68% of the response variance. The variance explained by each latent factor for the predictor and response datasets can be viewed separately. **(C)** Randomization Test Results: After shuffling the log2 fold changes, disrupting their correlations with input spectra, *MetaboLiteLearner*’s error with shuffled data was consistently higher than with the original dataset, confirming its ability to identify genuine relationships between metabolite spectra and abundance changes in rewired cells.

The model with *N*_*opt*_=5, when trained on the entire MLOD, optimally adjusted its loadings to maximize covariance between the projections of both inputs (the spectra) and outputs (log2 fold changes) onto the 5-dimensional latent space (**Fig. 3B**). Unlike unsupervised methods such as principal component analysis (PCA), which primarily focuses on variance within individual datasets, the PLSR emphasizes joint variance optimization (15). The resultant transformations into the latent space accounted for 32% of the predictor variance and 68% of the response variance. The variance explained by each latent factor for the predictor and response can be analyzed separately (**Fig. 3B’** and **Fig. 3B”**, respectively).

Next, we performed a randomization test to determine our model’s ability to capture biologically relevant patterns, not statistical artifacts. By shuffling the log2 fold changes, we retained internal correlations but disrupted correlations with input spectra. Notably, *MetaboLiteLearner*’s error, when trained with these shuffled data, was consistently worse than with the original dataset (**Fig. 3C**). This reaffirms our model’s capability to discern significant links between metabolite spectra and abundance shifts in rewired cells.

### *MetaboLiteLearner* Reveals Metabolic Changes in Metastatic Breast Cancer Cells

The optimal model with 5 latent components (*N*_*opt*_=5) trained on the entire MLOD can transform a spectrum (a 500-dimensional vector) into log2 fold changes for brain- and lung-homing cells (a 2-dimensional vector). This transformation can be visualized using a biplot, with each m/z ionic fragment represented by a vector (**Fig. 1A**). The biplot shows that specific fragments, such as m/z=104 which arises from the EI fragmentation of TMS derivatives of amino acids (14), correlate with increased levels in both cell types. In contrast, fragments like m/z=306, associated with EI-fragmentation of certain TMS-derivatized sugars (14), relate to decreased levels in organ-homing compared to parental cells.

Our previous study used an extensive panel of metabolites measured in targeted mode complemented with stable isotope tracing to analyze these cells (13). In that study, lung-homing cells displayed a pronounced Warburg effect, underscored by a high lactate dehydrogenase (LDH) to pyruvate dehydrogenase (PDH) expression ratio, a potential biomarker for lung metastasis.

To determine whether *MetaboLiteLearner* was discerning patterns of metabolic rewiring consistent with our previous study (13), we analyzed a set of 263 EI-fragmentation spectra for TMS-derivatized metabolites (18) which we grouped into seven categories from the KEGG database of compounds with biological roles (19). We fed the spectra into our trained model to predict fold changes. While most metabolites showed concurrent changes in both cell types, specific carbohydrates, and nucleic acids increased only in lung-homing cells (**Fig. 1A**).

The 5-latent-component model explains 67.7% of the variance in log2 fold changes (**Fig. 3B”**). This indicates that the model fits the data reasonably well. But beyond the quality of the fit, one of PLSR’s strengths lies in the possibility of dissecting the model’s coefficients to shed light on the molecular shifts underpinning cellular adaptations.

Components 1 and 3 showcase metabolites with decreased levels in both cell types, indicating overlapping metabolic shifts (**Fig. 4B**). Components 2 and 5 indicate metabolites with increased levels in both cell types. Component 4 highlights differences between the cell types—some metabolites decrease in brain-homing but increase in lung-homing cells. Component 1, which explains 27% of the response variance, captures a common trend—reduced levels in both cell types. Evaluating spectra from various compound classes, most amino acids follow this trend, whereas carbohydrates deviate from it (**Fig. 4C**). Analyzing all latent components (**Fig. 4D-G**) revealed component 4’s unique role. Despite accounting for just 4.9% of the response variance, it distinctly captures the variation between brain- and lung-homing cells. Carbohydrates and deoxyribonucleosides dominate this component, indicating potential metabolic shifts between the two cell types. These observed metabolic changes could reflect adaptations to the unique environments of the brain and lungs. With its rich blood supply, the lung might favor carbohydrate-utilizing cells. Conversely, the elevated deoxyribonucleosides in lung-homing cells could suggest more robust DNA repair mechanisms than in brain-homing cells. These findings align with our previous work (13) and suggest potential pathways for further exploration.

**Figure 4.**
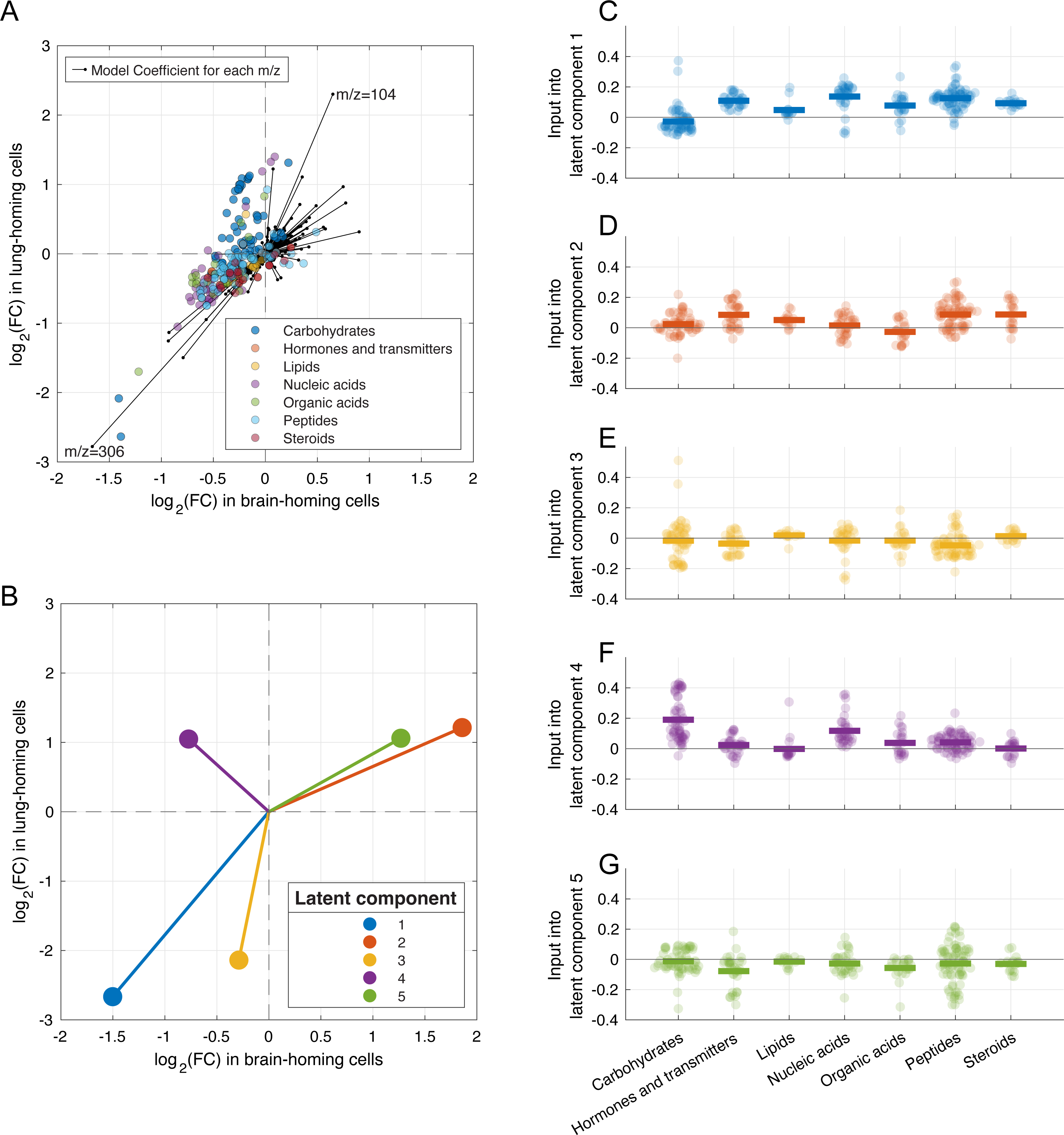
Interpretation of the Model for Metastatic Breast Cancer Cells. **(A)** Biplot representation of the m/z ionic fragment vectors. Specific fragments, such as m/z=104, are associated with increased levels in both cell types in contrast to fragments like m/z=306 which are associated with decreased levels. **(B)** The proportional contribution of the five latent components to the variance in log2 fold changes, with components 1 and 3 indicating overlapping metabolic shifts with decreased levels in both cell types, components 2 and 5 showcasing increases in both, and component 4 underscoring the divergence between the two cell types. **(C)** In component 1, accounting for 27% of the response variance, most amino acids follow the trend of reduced levels in both cell types, while carbohydrates differ. **(D-G)** Among the latent components, we see the distinctive role of component 4 which is dominated by carbohydrates and deoxyribonucleosides and highlights potential metabolic variances between brain- and lung-homing cells.

## Discussion

Here, we presented the capabilities of *MetaboLiteLearner*, a supervised learning approach to study the rewiring of intracellular metabolomes. Through computational experiments, we have shown that *MetaboLiteLearner* can discern patterns even within the confines of a relatively small dataset comprising 153 unidentified metabolites significantly altered in organ-homing cells compared to their parental cells, without prior knowledge of metabolic networks.

A defining feature of *MetaboLiteLearner*’s is its use of PLSR—a computationally lightweight and robust statistical algorithm, especially useful for smaller datasets. PLSR can handle internal correlations within predictors and responses, which arise due to shared ion fragments produced by naturally abundant isotopes and functional groups shared among certain metabolite classes and the universally witnessed metabolic shifts in disseminated cells, as noted in previous work (13). PLSR handles these internal correlations by generating a reduced-dimensional latent space that maximizes the joint variance between predictors and responses. Here, the latent space represents the leading associations between a metabolite’s molecular features (the ions produced by EI fragmentation) and its abundance change in rewired cells. While the current implementation of PLSR in our model is efficient, there is potential for further refinement, such as using regularization and Laplacian constraints (20). Incorporating regularization might improve model interpretability. Meanwhile, Laplacian constraints could let the model incorporate existing knowledge from comprehensive spectral libraries and metabolic network models.

In this study, we also introduce the MLOD, an open dataset tailored for machine learning research focusing on the metabolic reconfigurations in cancer cells. The MLOD, enriched with meticulously captured spectra and corresponding abundance shifts, can be a standard for benchmarking future supervised learning endeavors in this domain. Our decision to use GC/MS spectra from TMS-derivatized samples within a specific interval was primarily driven by its alignment with prevalent practices (18). However, we recognize the potential benefits of integrating data from high-resolution instruments like time-of-flight (TOF) or Orbitrap. These devices could capture finer details, thus enhancing model predictions. Furthermore, harnessing data generated by tandem instruments, such as MS/MS or MS2, can open doors to more sophisticated computational strategies, as witnessed in works external to cancer research (21– 23).

Our dataset focused on log2 fold changes from unsynchronized intracellular metabolite concentrations. This has constraints, as the insights obtained mainly reflect consistent changes across a generalized cell population. However, integrating synchronized data acquisition at different cellular growth stages (24) or incorporating stable isotope tracing (25) can inject dynamic elements into future datasets, allowing our models to capture more nuanced metabolic variations. While this study centered on breast cancer, the potential uses of *MetaboLiteLearner* extend beyond: it may be applied to a diverse range of cell types, conditions, and even less homogeneous samples like tissues or tumors. In our case study, the response variable was bidimensional capturing metabolite abundance changes in brain- and lung-homing cells relative to their parental lineage. The progression towards incorporating additional dimensions and more comprehensive datasets could require more sophisticated computational approaches, like deep neural networks, enabling a broader and deeper exploration of metabolic alterations across different cell conditions and types.

This study showcases the potential synergy between machine learning and metabolomics. With evolving datasets and improving computational methods, we can make new strides in unraveling the intricacies of metabolic rewiring—a fundamental aspect of cellular adaptation.

## Methods

### Cell Culture, Metabolite Extraction and Derivatization

Cell lines were cultured in DMEM (Fisher 11965118) supplemented with 10% FBS, produced in the MSKCC media core facility, and 1% penicillin/streptomycin (Fisher 15140122). The cultivation conditions included a 37°C incubator with regulated humidity and a 5% CO2 atmosphere. Authenticated cell lines were procured from the Massague lab and were developed as delineated in previous studies (11, 12). Cells were subjected to metabolite extraction using 1 ml of ice-cold 80% methanol, followed by overnight storage at -80°C. Subsequent to this, the extracts underwent a drying process using an evaporator. Resuspension was achieved by incubation with shaking at 30°C for 2 hours in a solution containing 50 μl of 40 mg ml−1 methoxyamine hydrochloride in pyridine.

Derivatization was performed by adding 80 μl of N-methyl-N-(trimethylsilyl) trifluoroacetamide (with or without 1% TCMS from Thermo Fisher Scientific) and 70 μl of ethyl acetate (sourced from Sigma-Aldrich). This mixture was then incubated at 37°C for 30 minutes.

### Gas Chromatography/Mass Spectrometry (GC/MS) Analysis

Analytical procedures utilized the Agilent 7890A gas chromatograph paired with an Agilent 5977C mass selective detector. The gas chromatograph operated in splitless injection mode, maintaining a constant helium gas flow at 1 ml/min. The injection involved introducing 1 μl of the derivatized metabolites onto an HP-5ms column. The temperature of the gas chromatograph oven was systematically ramped from 60°C to 290°C over a 25-minute interval. Samples comprised four distinct types: blank media, parental cells, lung-homing cells, and brain-homing cells. Each type was cultured in triplicate groups over a span of three days, resulting in nine replicates for each sample category.

### Data Processing

GC/MS raw data, stored in the Agilent.D format, underwent processing to generate the MLOD dataset. These raw.D files were initially converted into CSV. By creating a “virtual bulk sample” from this CSV data, we could detect and extract spectra from the total ion chromatogram (TIC). With the aid of the Matlab function *mspeaks*, peaks associated with different metabolites were identified within the TIC, followed by the extraction of their spectra. An integration process then yielded a peak area table. To maintain the relevance of the dataset, compounds that did not exhibit significant differences compared to blank media samples, as determined using ANOVA, were removed. Following this filtering process, the refined data formed the MLOD dataset, which has 153 unique spectra labeled with abundance alterations, represented as log2 fold changes for both brain-homing and lung-homing cells.

### Machine Learning

Supervised learning attempts to learn from training data containing inputs matched to their correct outputs, a model *Y = f(X)* that can generalize to a new dataset (15). During the learning phase, the model is a “student” given a set *X*_*train*_, *Y*_*train*_ of training. For each example *x*_*i*_ in the training data set *X*_*train*_ the model presents the answer *f(x*_*i*_ *) =ŷ*_*i*_. The supervisor or “teacher” compares *ŷ*_*i*_ with the correct answer, *y*_*i*_, and gives an error associated with the “student’s” answer. The examples’ errors are used to calculate a loss, *L(y*_*i*_, *ŷ)*, and the model learns by adjusting its parameters to minimize the loss. After the training, the model is presented with a test dataset *(X*_*teSt*_) containing a new set of examples. The model’s accuracy in predicting the output for the new inputs shown is evaluated by comparing these predictions *Ŷ*_*teSt*_ with the actual data, *Y*_*teSt*_.

*MetaboLiteLearner* learns a linear function *f* that takes a feature vector representation of a metabolite as input. It predicts, as output, how the intracellular level of that metabolite is impacted by metabolic rewiring. Each metabolite is represented by an input vector *x*_*i*_, a *p*-dimensional array of the features of metabolite *i*. In our case, *x*_*i*_ is 550-dimensional, representing the abundances of ionic fragments of metabolite *i* of sizes between 50 and 600 mass-to-charge ratio units (m/z) binned at unit intervals. This vector is obtained from the electron ionization (EI) mass spectrum. The EI-spectrum is a mass-to-charge histogram of the ion fragments produced when a metabolite undergoes breakdown via electron ionization. It offers a direct insight into the metabolite’s structure—and potentially its function—without needing to identify that metabolite first. The output vector *y*_*i*_ is two-dimensional, representing the log_2_ fold-change of that metabolite in brain-homing and lung-homing cells compared to the parental lineage.

*MetaboLiteLearner*’s learning algorithm is the partial least squares regression (PLSR) (16). This algorithm has similarities with artificial neural networks (17) but is more stable and less computationally demanding to run. We used the MATLAB implementation of PLSR encoded in function *plsregress*. Leave-one-out cross-validation was used to determine the optimal number of latent components, *N*_*opt*_. To ensure the robustness of the analytical outcomes, a shuffling test was administered, wherein the order of observed data was randomly rearranged a thousand times. Augmenting the biological context of the study, a comprehensive table detailing compounds with biological roles was sourced from the Kyoto Encyclopedia of Genes and Genomes (KEGG) (19). The spectra for TMS-derivatized compounds were obtained from the Fiehn Library (18). In conjunction with the KEGG table, these spectra were input into the *MetaboLiteLearner* model to compute their fold changes in brain- and lung-homing cells and the values for the five latent components.

## Data Availability

The raw GC/MS data supporting the findings of this study are available in Zenodo (26).

## Code Availability

The data processing and analysis code is available on GitHub (27).

## Acknowledgments

This work was supported by the National Institutes of Health (NIH) grant R01 CA266068 “Mathematical modeling of metabolism rewiring in cancer eco-evolution and metastasis tropism” to J.B.X. Acknowledgments to Chen Liao, Deepti Mathur, Heesoo Seong, Julia Simundza and Carlos Carmona Fontaine for their comments on the manuscript.

